# Chain-Length-Dependent Interactions of Polyethylene Nanoplastics with Human Catalase reveals Size based Nanoplastic Toxicity

**DOI:** 10.64898/2026.10.08.757803

**Authors:** Shristi Tanaya, Soumya Lipsa Rath

## Abstract

Nanoplastic contamination is increasingly recognized as a biochemical stressor, yet the molecular determinants governing how polymer fragment size dictates protein-level interactions remain poorly understood. Here, we combine molecular docking and all-atom molecular dynamics simulations to characterize the interaction of polyethylene (PE) oligomers of increasing length (n = 3, 5, 7, and 14 repeat units) with human catalase, a tetrameric heme enzyme central to cellular antioxidant defense. Single-chain simulations reveal that short PE fragments (3-, 5-, and 7-mer) migrate from an initial domain-connector pose toward a heme-proximal, previously characterized inhibitor-binding site, whereas the 14-mer remains kinetically confined to a peripheral surface site, implicating short-chain PE as a potential catalase inhibitor. In multichain corona systems, backbone dynamics show a non-monotonic, chain-length-dependent destabilization, with the 3- and 5-mer inducing the greatest structural perturbation and the 14-mer best preserving native-like dynamics despite forming an extensive, percolating surface network across all four subunits. MM-PBSA free energy decomposition identifies van der Waals interactions as the dominant thermodynamic driver of binding, while electrostatic surface mapping indicates that cationic surface patches instead govern long-range spatial targeting of PE chains. These findings establish a framework linking nanoplastic fragment size to protein-level toxicity, with direct implications for assessing the biochemical risk of environmentally weathered polyethylene.

## 1. Introduction

Plastic pollution represents one of the most pervasive anthropogenic stressors on contemporary ecosystems, and within this broader crisis, nanoplastics, particles with at least one dimension below 1 μm, have emerged as a subject of particular concern^25,2^. Their small size confers distinct physicochemical properties relative to microplastics (1 μm–5 mm), including a substantially higher surface area-to-volume ratio, enhanced environmental mobility, and, critically, the capacity to breach biological barriers that are otherwise impermeable to larger particulates, including the blood–brain barrier and the placental interface^1,2^. Once internalized, nanoplastics are not confined to the site of entry, they translocate across epithelial tissues and disseminate systemically to distal organs, including the liver, kidneys, brain, and placenta, raising fundamental questions about their capacity to perturb organismal homeostasis^3^.

In vitro and in vivo studies indicates that nanoplastic internalization initiates a convergent set of toxicological responses, including oxidative stress, mitochondrial dysfunction, endoplasmic reticulum stress, chronic inflammation, genotoxicity, and dysregulation of programmed cell death pathways^1,3^. The magnitude of these effects is modulated by particle size, polymer composition, exposure concentration, and surface chemistry, with smaller particles generally exhibiting greater cellular uptake and bioactivity^1,26^. Emerging evidence further suggests that nanoplastics may function as vectors for co-contaminants and disrupt inter-organ signaling, thereby amplifying systemic toxicity beyond what would be predicted from any single mechanism in isolation^27,28^. Among these interconnected outcomes, oxidative stress has been most consistently reported through which nanoplastic-induced cellular injury propagates^4^.

Catalase, is a heme-containing enzyme of exceptional catalytic efficiency, decomposing H₂O₂ into water and molecular oxygen at rates approaching diffusion-limited kinetics, with individual enzyme molecules capable of processing millions of substrate molecules per second^29,30^. Catalase also constitutes a enzymatic antioxidant defense system, without which cellular ROS accumulation would proceed unchecked^5^. H₂O₂ is generated continuously as a byproduct of mitochondrial respiration, peroxisomal β-oxidation, and the activity of numerous oxidase enzymes. While low concentrations of H₂O₂ contribute to physiological redox signaling, its accumulation drives Fenton chemistry and the generation of hydroxyl radicals, which indiscriminately damage lipids, proteins, and nucleic acids^31^. Catalase is located in the peroxisome and helps in buffering elevated ROS production beyond physiological set points due to different factors such as nanoplastic exposure^6,7^.

Structurally, catalase typically assembles as a homotetramer, with each subunit harboring a heme prosthetic group at the catalytic core^29^. The enzyme exhibits considerable stability across physiological ranges of pH and temperature, although its activity remains sensitive to modulation by inhibitors, activators, and the prevailing cellular redox state^5^. Impairment of catalase function has been implicated in a spectrum of pathological conditions, including diabetes, carcinogenesis, neurodegeneration, and accelerated aging, depicting its relevance to organismal health beyond its canonical antioxidant role^5,6,7^.

Given the consistency with which oxidative stress is reported across nanoplastic toxicity studies spanning unicellular algae, aquatic and terrestrial vertebrates, and human cell-based models, catalase activity has been increasingly employed as an index of nanoplastic-induced oxidative injury and of the organism’s compensatory capacity^8,32^. Suppressed catalase activity following exposure is generally interpreted as evidence that ROS generation has exceeded antioxidant defense capacity, often preceding more severe downstream pathology, whereas transient elevation may reflect an adaptive response of uncertain durability under prolonged or repeated exposure^6^. Monitoring these dynamics, in conjunction with complementary markers of lipid peroxidation, protein oxidation, and inflammatory activation, characterizing nanoplastic toxicity ^6,8^.

Environmental concentrations of nanoplastics, realistic human exposure levels, and dose–response relationships remain poorly understood, due to persistent analytical limitations such as:- no single technique currently provides comprehensive characterization of nanoplastic identity, morphology, and concentration within complex biological or environmental matrices, and standardized detection protocols have yet to be established^33,34^. Although catalase is a well-validated marker of oxidative stress in general, time-dependent response to nanoplastic exposure, as a function of particle size, polymer identity, and tissue type, remains inconsistently characterized ^6,9^.

Resolving these gaps carries both mechanistic and translational significance. The detection of nanoplastics in human blood, pulmonary tissue, and placental tissue has transformed concerns regarding chronic low-dose human exposure from a theoretical possibility into an empirically grounded priority for biomedical and environmental research^1,35^. Establishing the mechanistic pathways by which nanoplastics induce cellular injury, and at what exposure thresholds, is a prerequisite for evidence-based regulatory action.

Despite the growing evidence that nanoplastic contaminants can interact with and perturb biological macromolecules, the molecular determinants of these interactions, particularly how polymer chain length governs binding site selectivity, protein stability, and interfacial chemistry, remain poorly resolved at atomic resolution. Here, we systematically characterize the interaction of polyethylene (PE) oligomers of increasing length (n = 3, 5, 7, and 14 repeat units) with human catalase, a tetrameric heme enzyme central to cellular antioxidant defense, chosen as a physiologically relevant and mechanistically well-characterized target for nanoplastic exposure. We first examine single-chain PE binding to an isolated catalase subunit before extending this analysis to physiologically relevant multichain systems in which the full catalase tetramer is simultaneously exposed to many copies of a single PE chain length. Our analysis reveals a chain-length-dependent, non-monotonic relationship between PE size and protein structural stability, active-site inhibition potential, and PE self-association behavior. Together, these results establish a mechanistic framework for understanding how nanoplastic fragment size modulates protein-level toxicity, with direct implications for assessing the biochemical risk posed by environmentally weathered polyethylene degradation products.

## 2. Materials and Methods

### 2.1. System Preparation

The crystal structure of human catalase was retrieved from the RCSB Protein Data Bank (PDB ID: 7VD9). Because the deposited structure is a homotetramer, a single subunit (chain A) was extracted to serve as the protein model for all single-chain PE-binding simulations^10^. The N-terminal arm, which threads between neighboring subunits in the native tetramer and has no equivalent intra-subunit binding partner in the isolated monomer, was removed to yield a truncated, structurally self-consistent catalase construct suitable for standalone simulation. We also removed the bound NADPH from model.

Linear polyethylene (PE) oligomers of four chain lengths (3-, 5-, 7-, and 14-mer) were constructed as atomistic model nanoplastic fragments. The repeating –CH₂– unit of polyethylene was first encoded in SMILES (Simplified Molecular Input Line Entry System) notation, and oligomeric chains of the desired length were generated by iterative concatenation of this repeat unit, yielding the SMILES strings corresponding to PE3, PE5, PE7, and PE14. Each SMILES string was parsed into a molecular graph using the RDKit cheminformatics library in Python, and explicit hydrogen atoms were added to complete the all-atom connectivity^11,12^. Three-dimensional starting coordinates for each oligomer were then generated using the ETKDG (Experimental-Torsion Knowledge Distance Geometry) embedding algorithm implemented in RDKit, and the resulting geometries were energy-minimized using the Universal Force Field (UFF) to relax the initial embedding into a stable, low-energy conformation^12,13,14^. The optimized structures were exported in PDB format for use in subsequent docking and molecular dynamics steps. Prior to use, all PE PDB files were visually inspected in UCSF ChimeraX to confirm correct backbone connectivity, hydrogen placement, and overall geometric consistency; the verified structures served as the initial polymer models for all docking and Molecular Dynamics (MD) simulations described below^15^.

### 2.2. Molecular Docking and Corona Complex Construction

Molecular docking of each PE oligomer (PE3, PE5, PE7, PE14) against the prepared catalase subunit was performed using PatchDock. For each PE–catalase pair, multiple candidate docking poses were generated and ranked according to the PatchDock geometric shape-complementarity score^16^. To construct physiologically relevant, high-occupancy “corona” complexes representative of multivalent PE–protein exposure, overlapping and sterically clashing poses were identified and manually removed for each chain length, retaining only a non-redundant set of PE chains distributed across the catalase surface. This procedure yielded corona complexes containing 24 chains of PE3, 20 chains of PE5, 16 chains of PE7, and 7 chains of PE14 (Table 1). These non-overlapping, multichain PE–catalase complexes were used as the starting structures for all subsequent molecular dynamics simulations of the corona systems.

**Table 1.** Composition of the multichain PE–catalase corona complexes after removal of overlapping poses.

| PE oligomer | Chain length (monomers) | No. of chains in corona |
| --- | --- | --- |
| PE3 | 3 | 24 |
| PE5 | 5 | 20 |
| PE7 | 7 | 16 |
| PE14 | 14 | 7 |

### 2.3. Molecular Dynamics Simulations

All-atom molecular dynamics simulations of the PE–catalase corona complexes were performed using GROMACS version 2026.2 software, with the CHARMM36 force field applied to the protein and generated parameter from CGenFF for PE oligomers^17,18,19^. The non-overlapping, sterically resolved PE–catalase complexes generated as described in Section 2.3 served as the starting configurations for each simulation.

Each complex was placed in a rectangular simulation box and solvated explicitly using the TIP3P water model; the net system charge was neutralized by the addition of counterions. Each system was subjected to energy minimization using the steepest-descent algorithm to remove steric clashes and unfavorable contacts introduced during system construction. Equilibration was carried out in two stages: an initial NVT equilibration at 300 K with position restraints applied to both the protein and the PE chains, followed by NPT equilibration at 300 K and 1 bar using the V-rescale thermostat and the Parrinello–Rahman barostat to establish the correct system density and pressure^20^. Production simulations were subsequently performed for 200 ns in the NPT ensemble at 300 K and 1 bar. Long-range electrostatic interactions were treated using the particle mesh Ewald (PME) method, and van der Waals interactions were truncated at a cutoff distance of 1.2 nm. All bonds involving hydrogen atoms were constrained using the LINCS algorithm during production simulations, permitting an integration time step consistent with this constraint scheme^21^. All trajectories were analyzed using the built-in analysis tools of GROMACS and visualization was done using VMD (Visual Molecular Dynamic)^22^.

Binding free energies for the single-chain PE–catalase complexes (PE3, PE5, PE7, PE14) were calculated using the Molecular Mechanics Poisson–Boltzmann Surface Area (MM-PBSA) method, as implemented in gmx_MMPBSA. For each complex, snapshots were extracted at regular intervals from the equilibrated portion of the last 10 ns production trajectory. For each extracted snapshot, the binding free energy was decomposed as ΔGbind = ΔGmm + ΔGsolv − TΔS, where ΔGmm (reported as ΔGGAS) is the gas-phase molecular mechanics energy, comprising the van der Waals (ΔVDWAALS) and electrostatic (ΔEEL) interaction energy terms, and ΔGsolv (reported as ΔGSOLV) is the solvation free energy, comprising a polar contribution (ΔEPB), and a nonpolar contribution (ΔENPOLAR), estimated from the solvent-accessible surface area (SASA)^23, 24^.

## 3. Results and Discussion

Human catalase (PDB ID: 7VD9) is a homotetrameric heme enzyme in which each subunit adopts the canonical catalase fold, organized into four structurally discrete regions: an N-terminal arm (residues 5–70), a core β-barrel domain (residues 71–379), a wrapping loop (residues 380–438), and a C-terminal helical domain (residues 439–502) (Figure 1a). The N-terminal arm, although largely devoid of secondary structure beyond a single short α-helix, threads between neighboring subunits and thereby locks the tetramer together, consistent with a cooperative, interdigitated assembly pathway rather than simple docking of independently folded monomers. The catalytic core comprises an eight-stranded antiparallel β-barrel flanked by α-helices, which houses the heme b prosthetic group deep within the subunit interior, shielded from bulk solvent. The wrapping loop, though structurally irregular, mediates extensive interdomain and inter-subunit contacts, while the C-terminal helical domain contributes to oligomeric stability and links the buried active site to the protein surface.

**Figure 1.**
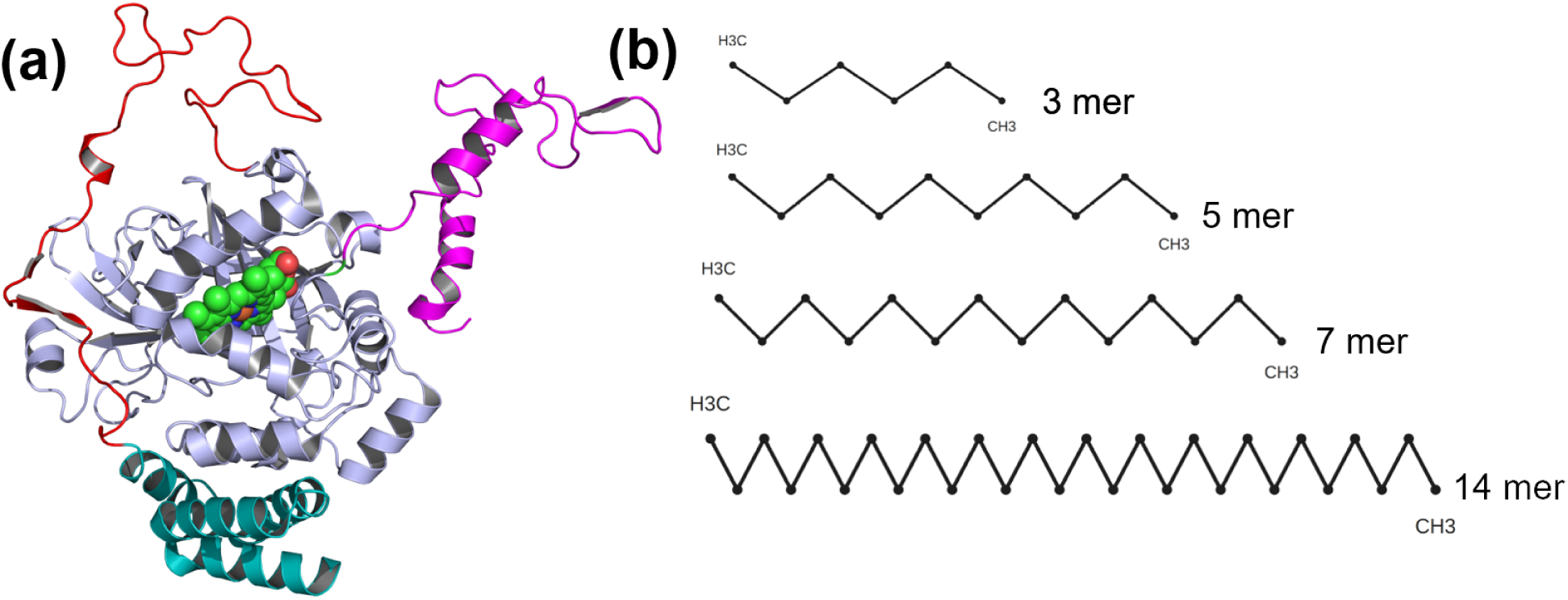
Structural overview of catalase and polyethylene (PE) oligomer models. (a) Cartoon representation of a single catalase subunit (chain A, PDB ID: 7VD9), color-coded by structural region: N-terminal arm (residues 5–70), core β-barrel domain (71–379), wrapping loop (380–438), and C-terminal helical domain (439–502); the heme *b* cofactor is shown in stick representation at the subunit core. (b) Atomistic models of the linear PE oligomers used in this study (3-, 5-, 7-, and 14-mer), generated by SMILES-based chain construction.

As a model nanoplastic system, we selected linear polyethylene (PE) oligomers of four lengths (n = 3, 5, 7, and 14 repeat units) (Figure 1b), which consist of repeating –CH₂– units linked into an all- carbon backbone. Short oligomeric PE chains provide a tractable means of probing how backbone length governs conformational behavior: at n = 3–5, chain-end effects dominate, rotation about the C–C bonds is only weakly hindered by neighboring segments, and the chain samples a broad ensemble of gauche and trans conformers with little tendency toward an extended geometry. At n = 7, cooperativity between adjacent torsional angles begins to favor extended all-trans stretches and short-range order emerges, although the chain remains too short to adopt a well-defined crystalline packing motif. By n = 14, the oligomer approaches bulk-like PE behavior, preferentially adopting the all-trans planar zig-zag backbone characteristic of the orthorhombic PE lattice, with CH₂ units spaced at the canonical ∼2.53 Å repeat along the chain axis. These oligomers are directly relevant to the study of nanoplastics, since environmental weathering of bulk PE proceeds through progressive oxidative and mechanical chain scission that reduces long polymer chains to short, low-molecular-weight fragments prior to their release as free nanoparticles or solubilized species. An oligomer series spanning n = 3–14 therefore serves as an atomistic proxy for these degradation intermediates, enabling us to probe how chain length governs physicochemical behavior across the transition from environmentally mobile fragment to bulk polymer. This is particularly relevant given that the electronic properties of PE, and by extension its potential for oxidative reactivity toward biomolecules, are known to vary systematically with oligomer size.

### A. Single-Chain PE Binding to Catalase

We first docked catalase with individual PE chains of each length (n = 3, 5, 7, 14) and subjected the top-scoring pose for each system to 200 ns of MD simulation in triplicate. Backbone Cα RMSD (Figure S1a) showed rapid initial relaxation in all systems, followed by convergence to a stable plateau of ∼0.5–0.7 nm after 50–100 ns, indicating that each complex reached a well-equilibrated conformational state within the simulated time window. Per-residue RMSF analysis (Figure S1b) showed that the protein core (residues 70–370 and 440–500) remained rigid across all replicates (RMSF < 0.2 nm), whereas the wrapping loop (residues 380–430) exhibited pronounced, reproducible flexibility, reaching RMSF values of up to ∼1.2 nm irrespective of PE chain length. Because this loop directly mediates inter-subunit contacts within the physiological tetramer, its intrinsic flexibility in the single-subunit simulations is consistent with a functional role in engaging the neighboring subunit interface, and, as discussed below, in accommodating bound PE.

Comparison of the initial and final (200 ns) PE poses (Figure 2) revealed marked chain-length-dependent binding behavior. The top-scoring poses for the 3-, 5-, and 7-mer chains initially localized to the domain connector (wrapping) loop region, whereas the 14-mer instead scored highest at the C-terminal α-helical domain. Over the course of the simulations, the shorter chains (3-, 5-, and 7-mer) proved highly mobile, repositioning independently in each replicate; their most frequently sampled site was adjacent to the heme, at the interface between the β-barrel and the domain connector loop. In contrast, the 14-mer remained stably bound at its initial C-terminal site in all three replicates, indicating that chain length imposes a kinetic barrier to large-scale repositioning, consistent with the greater number of simultaneous protein contacts available to a longer, more extended chain. This migratory behavior is notable in light of previously reported catalase inhibitor-binding sites: site 1, adjacent to the active-site heme, and site 2, near the domain connector loop ^37^. Although the best-scoring docked poses for the shorter oligomers were initially located near site 2, all three (3-, 5-, and 7-mer) consistently migrated toward site 1 over the course of the simulations, suggesting that site 1 is energetically more favorable for PE binding and raising the possibility that these short-chain PE fragments could act as catalase inhibitors in a manner analogous to previously characterized site 1-binding ligands. To quantify this preference, we calculated docking energies for the 3-, 5-, and 7-mer chains at both sites (Table S1). Binding affinities at the two sites were comparable overall, with a modest preference for site 1, consistent with its more buried, heme-proximal environment providing additional van der Waals contacts relative to site 2, which lies adjacent to the more solvent-exposed and conformationally flexible domain connector loop. Affinity at both sites increased with increasing PE chain length, reflecting the larger contact area available to longer oligomers.

**Figure 2.**
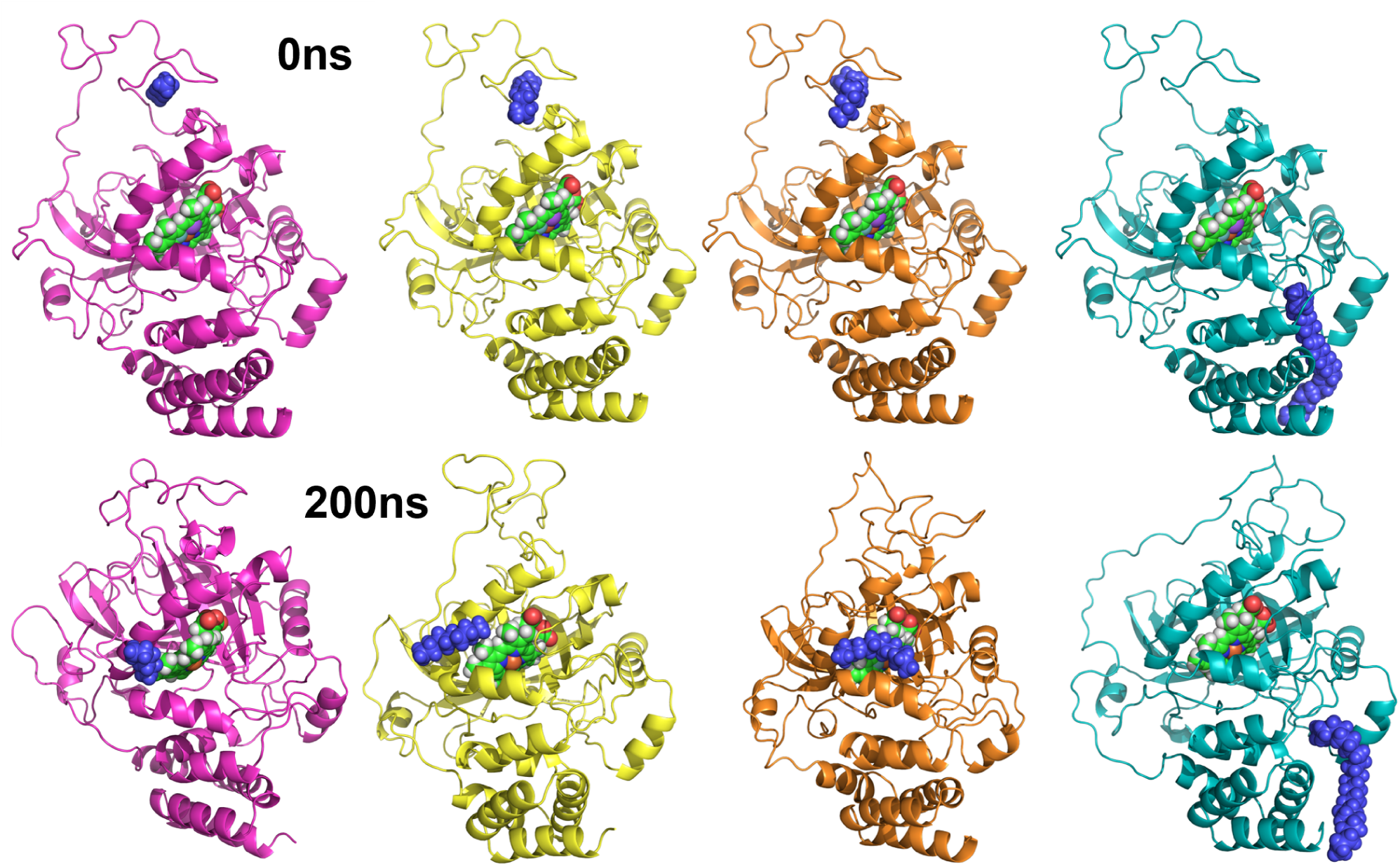
Chain-length-dependent binding site migration of single PE chains on catalase. Representative starting (top row) and 200 ns-equilibrated (bottom row) conformations of catalase in complex with a single bound PE chain: 3-mer (magenta), 5-mer (yellow), 7-mer (salmon), and 14- mer (cyan); PE monomer units are shown as spheres and the heme cofactor in stick representation. The 3-, 5-, and 7-mer chains reposition substantially over the trajectory, migrating from the initial domain-connector pose (site 2) toward the heme-proximal site (site 1), whereas the 14-mer remains stably bound at its initial C-terminal site throughout.

DSSP-based secondary structure analysis of PE-bound catalase relative to the wild-type (WT) apo structure revealed region-specific, chain-length-dependent perturbations. α-Helical content was selectively lost in the 240–260 region in the 7-mer complex, while β-sheet disruption at residues 165–180 occurred in both the 5-mer and 14-mer complexes, and helical loss near the C-terminus (452–470) was shared by the 7-mer and 14-mer complexes. The 5-mer complex exhibited the most extensive and widespread structural perturbation of the three variants examined, whereas the loop region spanning residues 368–440 remained conformationally stable in both WT and the 14-mer complex. Notably, all three PE-bound variants showed convergent structural alterations within residues 212–225, identifying this segment as a chain-length-independent hotspot of PE-induced conformational sensitivity in catalase.

MM-PBSA decomposition confirmed that binding free energy becomes progressively more favorable with increasing PE chain length (ΔTotal = −6.12, −8.89, −14.94, and −18.8 kcal/mol for the 3-, 5-, 7-, and 14-mer, respectively) (Table 2). The van der Waals interactions constitutes the dominant favorable contribution at all chain lengths and electrostatic (ΔEEL) contributes negligibly throughout. Notably, the 5-mer exhibited a disproportionately large polar desolvation penalty (ΔEPB = +11.55 kcal/mol) relative to its van der Waals contribution, yielding a less favorable net affinity than the 7-mer. Thuys, although comparable nonpolar packing exist among the 5-mer and 7- mer complexes, the greater structural destabilization observed for the 5-mer–catalase complex.

**Table 2.** MM-PBSA Binding Free Energy Decomposition for PE–Catalase Complexes as a Function of Chain Length.

| PE oligomer | $\Delta\text{VDWAALS}$ | $\Delta\text{EEL}$ | $\Delta\text{EPB}$ | $\Delta\text{ENPOLAR}$ | $\Delta\text{GGAS}$ | $\Delta\text{GSOLV}$ | $\Delta\text{Total}$ |
| --- | --- | --- | --- | --- | --- | --- | --- |
| PE3 | -7.27 | -0.1 | 2.48 | -1.23 | -7.37 | 1.25 | -6.12 +/- 0.96 |
| PE5 | -17.8 | -0.12 | 11.55 | -2.51 | -17.92 | 9.03 | -8.89 +/- 1.13 |
| PE7 | -17.64 | -0.33 | 5.35 | -2.33 | -17.96 | 3.02 | -14.94 +/- 1.33 |
| PE14 | -28.9 | -0.47 | 14.34 | -3.77 | -29.37 | 10.57 | -18.80 +/- 1.46 |

### B. Multichain PE Binding to Catalase

To capture more physiologically relevant, higher-occupancy binding scenarios, we extended this analysis to multichain systems in which the full catalase tetramer was simultaneously exposed to many copies of a single PE chain length. After docking and removal of overlapping poses, 24 chains of the 3-mer, 20 chains of the 5-mer, 16 chains of the 7-mer, and 7 chains of the 14-mer were retained. The number of independent, non-overlapping high-scoring poses accommodated on the tetramer surface were found to be decreasing with increasing chain length, as expected due to steric considerations. Each heme-bound, PE-decorated tetramer was simulated for 200 ns.

The Cα RMSD of the tetramer backbone (Figure 3a) revealed distinct conformational stability profiles across the WT and PE-bound variants (1PE3mer, 1PE5mer, 1PE7mer, 1PE14mer). WT catalase equilibrated rapidly within the first ∼20 ns and stabilized at 0.55–0.60 nm for the remainder of the trajectory, with minimal fluctuation, characteristic of a well-converged native fold. The 1PE14mer complex displayed the lowest overall RMSD of all five systems, plateauing near 0.45– 0.50 nm after an initial rise suggesting that binding of the longest PE chain does not destabilize the tetramer backbone and may confer a modest stabilizing effect, potentially through extended, multivalent surface contacts that restrict large-scale backbone motion (see below). In marked contrast, the 1PE3mer and 1PE5mer complexes exhibited substantially elevated RMSD, fluctuating persistently between 0.85 and 1.1 nm with pronounced, high-amplitude oscillations and no clear convergence to a stable plateau over the 200 ns trajectory, indicating sustained conformational drift from the native fold. This is consistent with the extensive secondary structure loss noted above for the 1PE5mer variant. The 1PE7mer complex showed intermediate behavior, rising from baseline to stabilize at 0.75–0.80 nm after ∼60–70 ns. Thus, the data reveal a non-monotonic, chain-length-dependent relationship between PE binding and catalase backbone stability: the shorter fragments (3-mer, 5-mer) induce the greatest deviation from the native conformation, the 7-mer produces intermediate destabilization, and the longest chain (14-mer) is best tolerated, most closely approximating native-like dynamics. This pattern indicates that PE-induced structural perturbation of catalase is not a simple function of nanoplastic size, but instead depends on chain-length-specific binding geometries and interfacial contact patterns.

**Figure 3.**
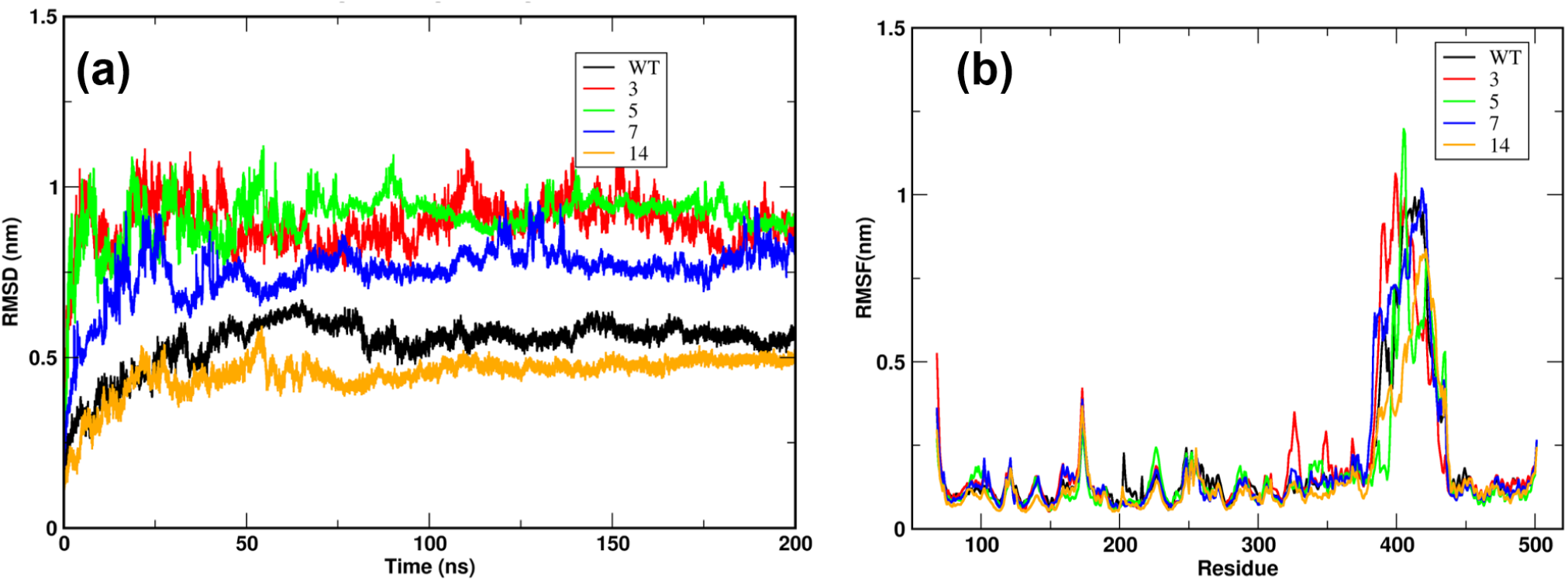
Backbone dynamics of multichain PE–catalase corona complexes. (a) Cα RMSD and (b) per-residue RMSF of the catalase tetramer backbone over 200 ns, for the apo wild-type (WT, black) and PE-bound corona complexes (1PE3mer, red; 1PE5mer, green; 1PE7mer, blue; 1PE14mer, magenta). WT and 1PE14mer equilibrate to the lowest, most stable RMSD plateaus, whereas 1PE3mer and 1PE5mer show persistently elevated, non-converging RMSD. RMSF traces identify the wrapping loop (residues 380–430) as a shared flexibility hotspot across all systems, with the largest-amplitude fluctuations observed for 1PE3mer and 1PE5mer.

Per-residue RMSF (Figure 3b) showed that all five systems exhibited closely overlapping, low-amplitude fluctuations (generally < 0.25 nm) across the majority of the β-barrel and helical scaffold, indicating that the core globular fold remains largely rigid and unperturbed by PE binding irrespective of chain length; PE-induced destabilization, where present, is therefore localized rather than distributed uniformly across the protein. The most prominent feature in all five traces is a pronounced flexibility hotspot spanning residues 380–430 (RMSF 0.6–1.2 nm), consistent with the solvent-exposed, partially unstructured wrapping loop identified in the single-chain simulations. Within this region, the 1PE3mer and 1PE5mer complexes showed the highest peak fluctuations (approaching 1.0–1.2 nm), exceeding both WT and the 1PE7mer and 1PE14mer complexes, indicating that the shorter PE fragments selectively amplify local loop destabilization in this segment. The 1PE14mer trace, by contrast, showed a comparatively dampened and right-shifted peak in this region relative to WT, implying that the longest PE chain partially restricts loop motion, consistent with its lower overall RMSD. Two secondary, more moderate flexibility peaks were also evident: one near residue 70, most prominent in the 1PE3mer complex, (∼0.5 nm) and another near residue 170, found in all the systems at comparable magnitude, ∼0.35–0.4 nm. The selective appearance of these additional peaks in the 1PE3mer trace suggests that the shortest oligomer engages a distinct, locally more disruptive binding mode at these sites relative to the longer chains.

Comparison of the initial and 200 ns PE conformers revealed clear chain-length-dependent differences in surface localization (Figure 4). The shorter chains, specifically the 3-mer and 5-mer, readily migrated into the interior of the catalase subunit over the simulated timescale, whereas the 7-mer chains remained bound within deeper surface grooves without fully penetrating the subunit core. The 14-mer, by contrast, did not traverse into the protein interior at any point and remained confined to the exterior surface throughout the trajectory, a restriction we attribute to its greater chain length precluding access through the same surface openings available to the shorter, more compact oligomers.

**Figure 4.**
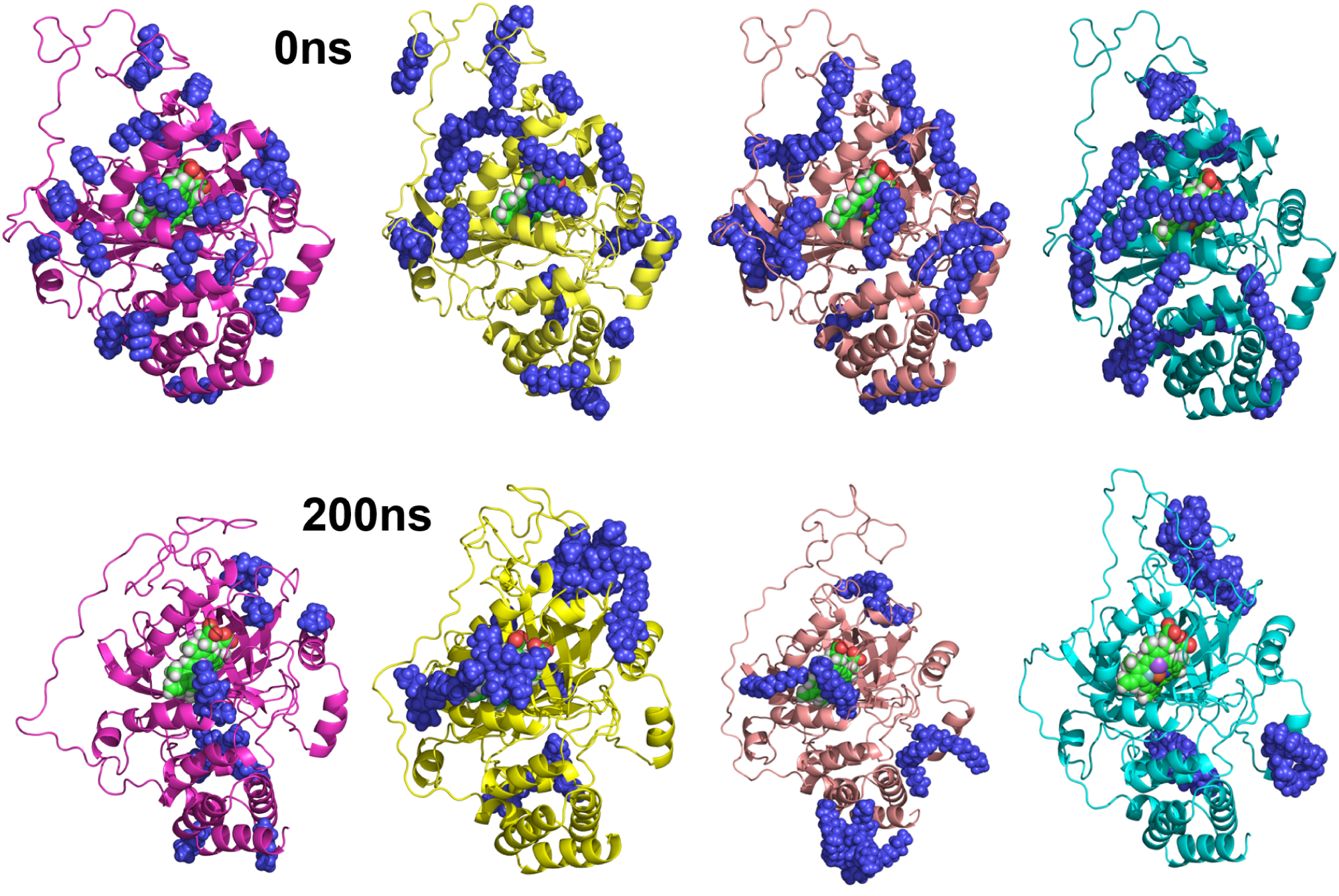
Chain-length-dependent surface localization of PE chains in multichain corona simulations. Representative initial (t = 0) and final (200 ns) conformations of PE chains bound to the catalase tetramer at each chain length, illustrating that the 3- and 5-mer chains migrate into the subunit interior over the simulated trajectory, the 7-mer remains bound within deep surface grooves without full penetration, and the 14-mer remains confined to the exterior surface throughout.

PE chains also formed compact, localized clusters at the protein surface, and cluster-number analysis (Figure 5; Table 3) showed that the absolute cluster count decreased monotonically with increasing chain length. However, because the number of PE chains present differed by system (24, 20, 16, and 7 chains for the 3-, 5-, 7-, and 14-mer systems, respectively), we additionally normalized the cluster count by the total chain population of each system (N/N₀) to isolate the intrinsic, per-chain propensity for self-association from the trivial effect of chain count on absolute cluster number (Table 3). This normalization revealed a distinctly non-monotonic trend: the intermediate-length chains (5-mer and 7-mer; N/N₀ ≈ 0.34–0.35) showed the greatest clustering efficiency per chain, whereas both the 3-mer and, notably, the 14-mer showed comparatively less efficient consolidation (N/N₀ ≈ 0.46 and 0.57, respectively). This indicates that the low absolute cluster count of the 14-mer system is driven substantially by its smaller total chain population rather than by an intrinsically higher per-chain aggregation propensity, and that intermediate chain lengths are in fact the most efficient at self-association on the catalase surface.

**Figure 5.**
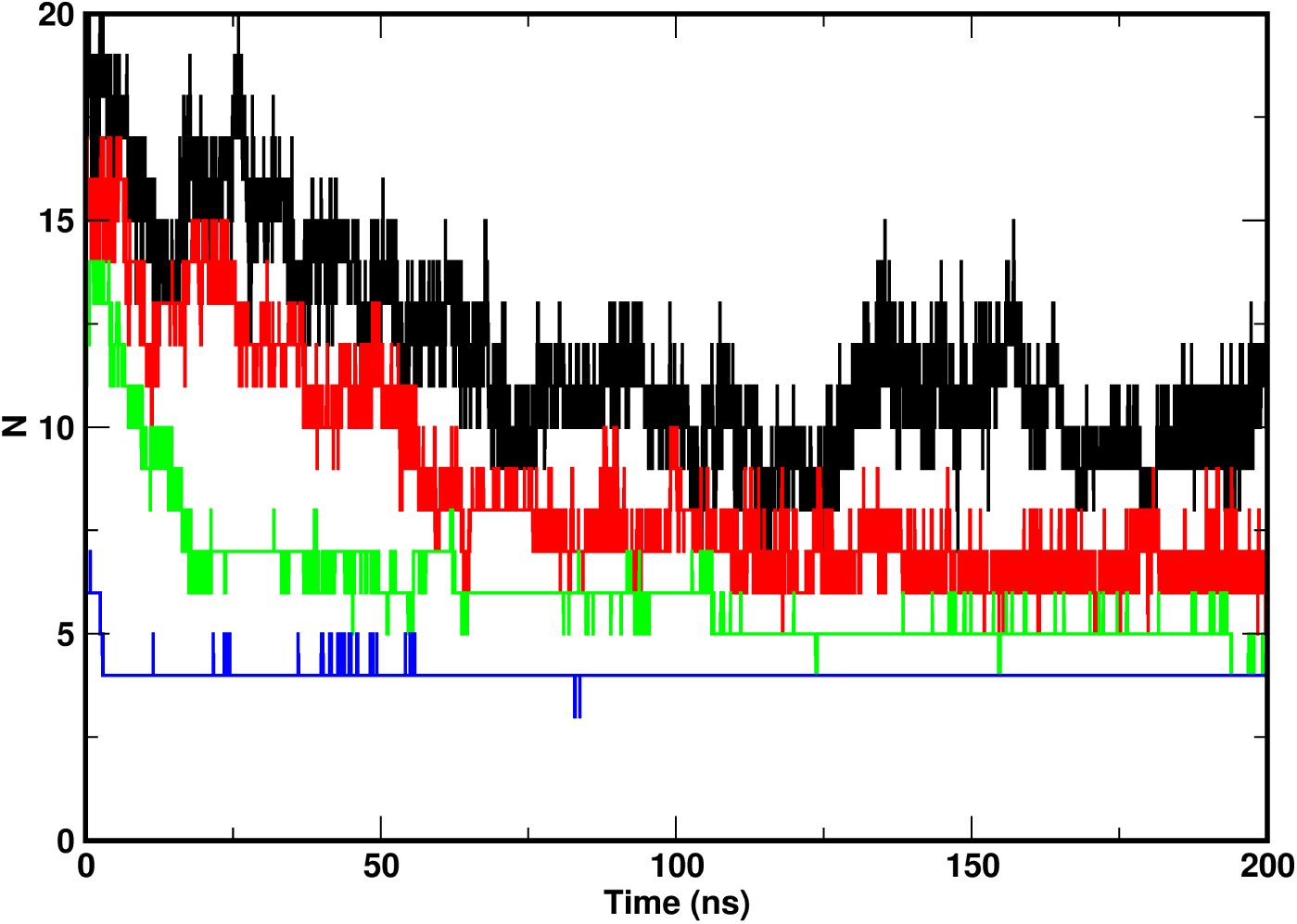
Time evolution of PE cluster number during multichain corona simulations. Number of distinct PE clusters (N) as a function of simulation time (0–200 ns) for the 3-mer (black), 5-mer (red), 7-mer (green), and 14-mer (blue) corona systems. All systems show a decline in cluster number from an initially dispersed state, converging to a stable plateau within 50–75 ns (∼10 ns for the 14-mer).

**Table 3.** Chain-Population-Normalized Cluster Analysis of PE Self-Association on the Catalase Surface.

| PE oligomer | Total chains ($N_0$) | Initial N ($t=0$) | Plateau N (~100–200 ns) | $N/N_0$ (plateau) | % reduction in cluster count |
| --- | --- | --- | --- | --- | --- |
| PE3 | 24 | ~19 | ~11 | 0.46 | ~42% |
| PE5 | 20 | ~16 | ~7 | 0.35 | ~56% |
| PE7 | 16 | ~13 | ~5.5 | 0.34 | ~58% |
| PE14 | 7 | ~6 | ~4 | 0.57 | ~33% |

To probe the physicochemical basis of PE recruitment to the catalase surface, we computed the APBS electrostatic potential surface of the tetramer (±3 kT/e) and mapped the positions of bound PE chains onto it (Figure 6). PE chains preferentially adsorbed at discrete electropositive surface patches rather than distributing uniformly across the protein: a single dominant cationic groove on the front face of the tetramer recruited the majority of bound chains, while two additional, lower-occupancy electropositive sites were identified on the reverse face. This spatial correspondence between PE occupancy and locally concentrated positive surface potential is consistent with a role for the protein’s electrostatic surface topology in biasing the diffusional approach and initial localization of PE chains toward a small number of preferred sites, rather than distributing adsorption stochastically across the available surface. This is in line with the MM-PBSA binding free energy decomposition above, which showed that the short-range energetics of the bound complex are dominated by van der Waals rather than electrostatic contributions. Together, the two analyses support a picture in which electrostatic surface topology governs long-range targeting, while hydrophobic packing governs short-range thermodynamic stabilization once PE chains reach the protein surface.

**Figure 6.**
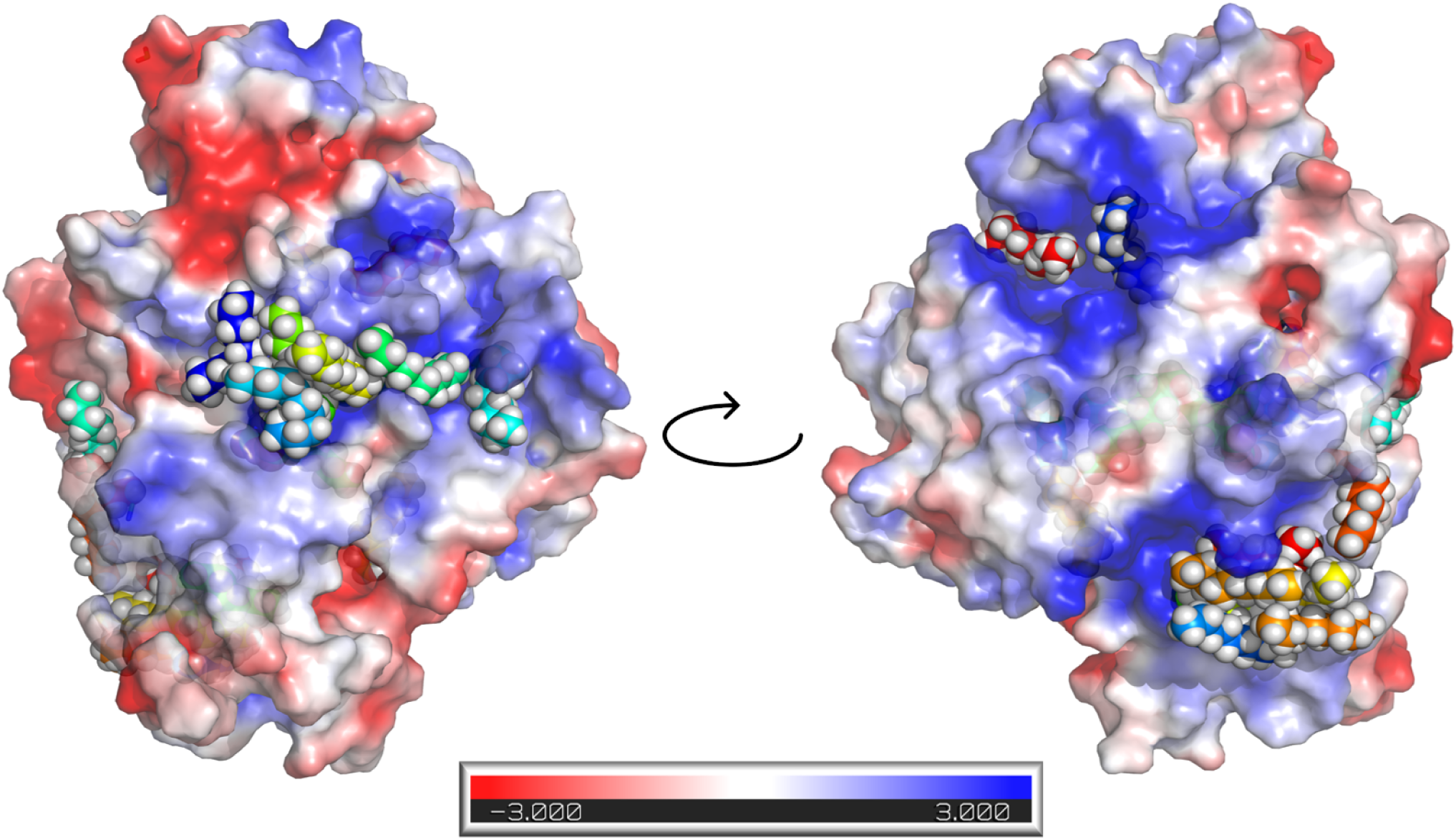
Electrostatic basis of PE adsorption on the catalase surface. APBS-calculated electrostatic potential surface of the catalase tetramer (red, −3 kT/e; white, neutral; blue, +3 kT/e), shown in front (left) and back (right, rotated 180°) views, with bound PE chains overlaid as space-filling spheres (colors denote separate PE chains). PE chains preferentially localize at electropositive (blue) surface patches, including a dominant cationic groove on the front face and two lower-occupancy sites on the reverse face.

### C. Length-Dependent Surface Coverage on the Catalase Tetramer

Visualization of PE occupancy across the full catalase tetramer as a function of chain length (Figure 7a–d) revealed a clear transition in adsorption mode. The shortest chains (3-mer) formed discrete, spatially isolated clusters confined to a small number of preferred surface sites and many units getting adsorbed into the protein tetramer. As chain length increased (5-mer, 7-mer), PE density and adsorption was found to increase. Additionally, the PE chains began to visibly bridge across subunit boundaries, indicating that chains of sufficient length can simultaneously engage multiple subunits rather than remaining confined to a single local site. At the longest chain length (14-mer), PE coverage increased dramatically, forming a dense, interconnected network that wrapped across nearly the entire solvent-accessible surface of the tetramer, engaging all four subunits simultaneously. This length-dependent transition from discrete, localized clusters to an extensively bridged, percolating surface network is consistent with the electrostatically nucleated, cooperative adsorption mechanism inferred from the electrostatic surface mapping and cluster-number analyses above: once nucleated at a cationic hotspot, sufficiently long chains possess the contour length required to propagate laterally across adjacent subunit surfaces and interconnect otherwise spatially separate binding sites.

**Figure 7.**
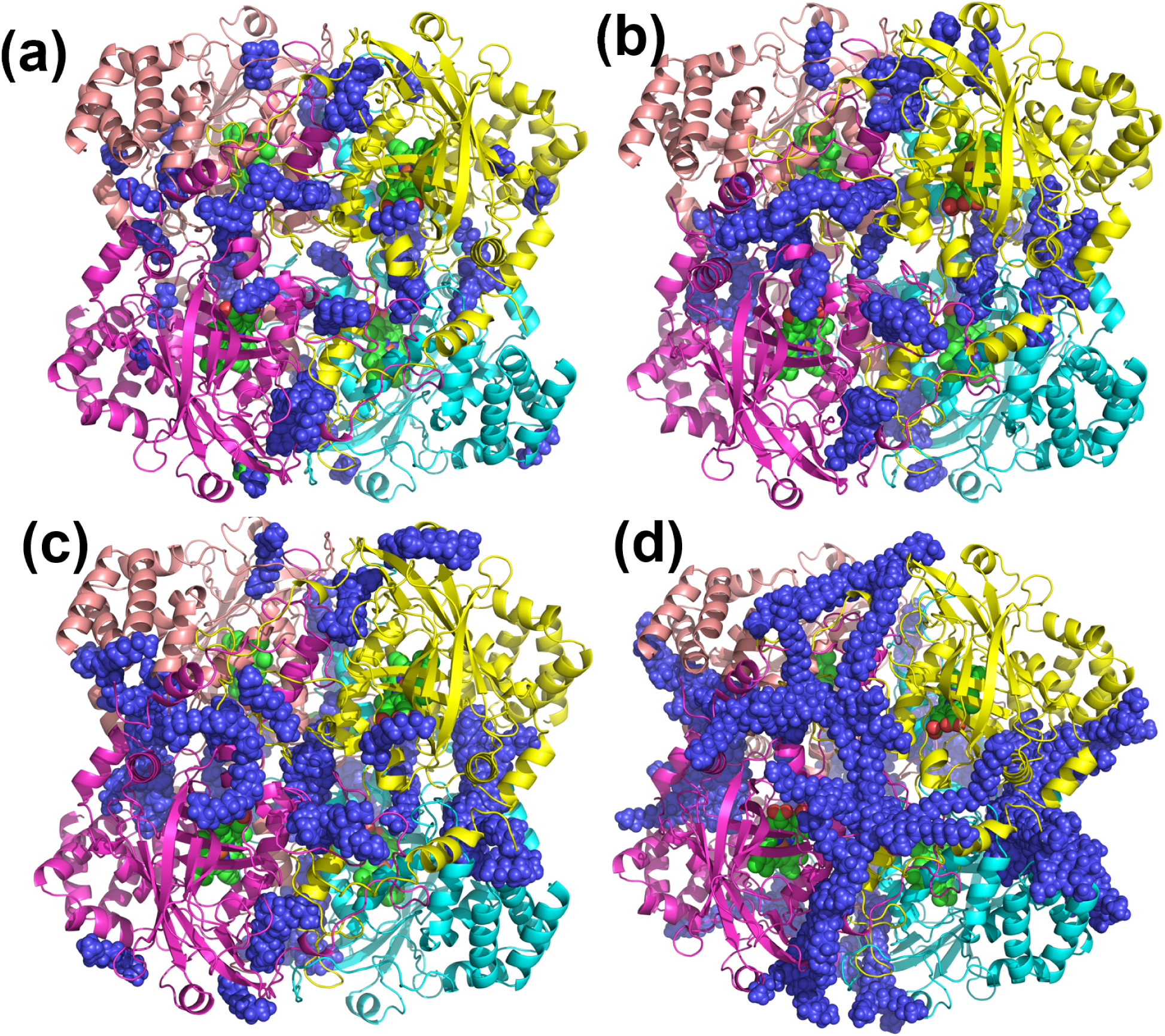
Length-dependent PE surface coverage across the catalase tetramer. Cartoon representation of the catalase tetramer (subunits colored salmon, magenta, yellow, and cyan; heme cofactors shown as sticks) in complex with bound PE chains (blue spheres) of increasing length: (a) 3-mer, (b) 5-mer, (c) 7-mer, and (d) 14-mer. Coverage progresses from discrete, spatially isolated clusters confined to a small number of preferred sites (a) to an extensive, interconnected network bridging all four subunits (d).

To characterize the chemical nature of the PE contact shell, we classified all catalase residues found in contact with bound PE chains, pooled across all four chain lengths, into acidic, basic, hydrophilic, and hydrophobic categories (Figure 8). This analysis revealed a strong compositional bias toward hydrophobic residues, which accounted for approximately 62% of all protein–PE contacts, greater than basic (∼16%), hydrophilic (∼16%) residues, and acidic residues (∼7%). This distribution indicates that, while long-range electrostatic complementarity governs the initial recruitment of PE chains to cationic surface patches (Figure 6), the resulting complex is predominantly stabilized at short range by hydrophobic packing rather than by persistent salt-bridge or hydrogen-bonding interactions. This two-stage picture, electrostatically driven long-range recruitment followed by hydrophobically dominated short-range stabilization, parallels the adsorption behavior reported for other protein–polymer and protein–nanoparticle corona systems, and, because the hydrophobic-contact dominance observed here holds when pooled across all four oligomer lengths, appears to be a chain-length-independent feature of the PE–catalase interface rather than a property specific to any single PE size.

**Figure 8.**
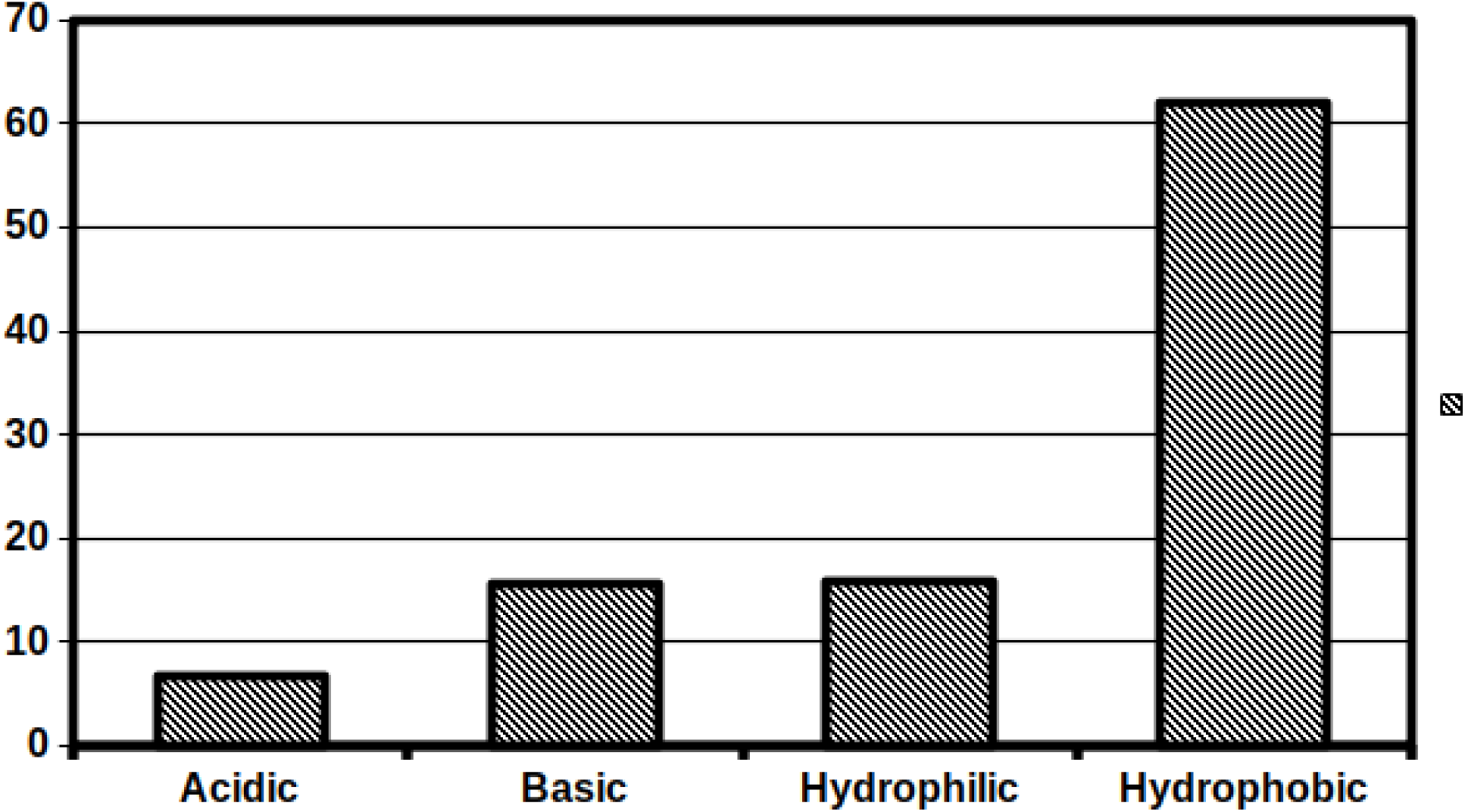
Residue-type composition of the PE–catalase contact shell. Percentage distribution of catalase residues in direct contact with bound PE chains, classified as acidic, basic, hydrophilic, or hydrophobic, pooled across all four PE chain lengths. Hydrophobic residues account for the majority of contacts (∼62%), substantially exceeding basic (∼16%), hydrophilic (∼16%), and acidic (∼7%) residue contributions.

## Conclusion

This work provides an atomistic picture of how polyethylene (PE) nanoplastic fragments of increasing chain length (n = 3, 5, 7, 14) interact with human catalase, both as single bound chains and as multichain surface assemblies, and links these interactions to measurable perturbations of protein structure and dynamics. Across both the single- and multichain systems, PE chain length emerged as the dominant variable governing binding mode, protein stability, and interfacial chemistry, though its influence was consistently non-monotonic rather than simply scaling with chain size.

At the single-chain level, the shorter oligomers (3-, 5-, and 7-mer) were conformationally mobile and migrated from an initially favorable domain-connector site (site 2) toward the heme-proximal inhibitor-binding pocket (site 1), a previously characterized site of catalase inhibition, whereas the 14-mer remained kinetically trapped at its initial C-terminal binding site throughout the simulated trajectory. This migratory behavior, combined with the favorable binding energies calculated for site 1, suggests that short-chain PE fragments in particular have the potential to act as catalase inhibitors, a mechanistic hypothesis that merits direct enzymatic validation.

At the multichain level, backbone RMSD and RMSF analyses revealed that the shorter fragments (3-mer, 5-mer) are the most structurally destabilizing, inducing sustained conformational drift and amplified flexibility at the wrapping loop (residues 380–430), while the 14-mer was the best tolerated, closely approximating native-like dynamics and, in the case of the longest chain, was restricted entirely to the exterior protein surface rather than penetrating into the subunit interior as the shorter chains did. Cluster-number analysis, when normalized by the total chain population of each system, further showed that clustering efficiency is not a monotonic function of chain length: intermediate-length chains (5-mer, 7-mer) self-associate most efficiently on a per-chain basis, while the 14-mer’s apparently low cluster count is largely an artifact of its smaller total chain population rather than an intrinsically greater aggregation propensity.

Electrostatic surface mapping and residue-level contact analysis together indicate a two-stage adsorption mechanism in which PE chains are first recruited to discrete, electropositive surface patches by long-range Coulombic interactions, and are subsequently stabilized at short range predominantly through hydrophobic packing (∼62% of all contacts), a feature that was consistent across all four chain lengths examined. Visualization of PE occupancy across the full catalase tetramer showed that this electrostatically nucleated adsorption manifests differently depending on chain length: short chains remain as discrete, spatially isolated clusters confined to a small number of preferred sites, whereas the longest chains grow laterally from these nucleation points into an extensive, percolating network that bridges all four subunits of the tetramer.

Collectively, these findings demonstrate that PE nanoplastic fragments perturb catalase through chain-length-dependent, site-specific mechanisms rather than a single, size-scaling mode of interaction, with the shortest oligomers posing the greatest risk of active-site engagement and structural destabilization, and the longest oligomers instead favoring extensive, cooperative surface coverage without penetrating or substantially destabilizing the native fold. Given catalase’s central role in cellular antioxidant defense, these results suggest that the smaller, more mobile PE degradation products generated during environmental weathering may represent a disproportionately greater biochemical hazard than their parent bulk polymer. This also presents the importance of oligomer size as a determinant of nanoplastic–protein interaction and downstream toxicological risk. Future work combining these computational predictions with direct enzymatic activity assays and mass-spectrometry-based interaction mapping will be important to establish whether the inhibitory and destabilizing effects predicted here translate into measurable losses of catalase function in vitro and in vivo.

## Acknowledgements

Authors thank the National Institute of Technology, Warangal, for providing computational facility and MHRD for the research scholar scholarship.

