## Supplementary Information for "Chain-Length-Dependent Interactions of Polyethylene Nanoplastics with Human Catalase reveals Size based Nanoplastic Toxicity"

**Table S1. Comparision of docking energy of nanoplastics at binding site 1 and 2****

| Reference ligand binding site 1 ^37^ | -7.8 and -7.7 kcal/mol |
| --- | --- |
| Reference ligand binding site 2 ^37^ | -7.8 and -7.5 kcal/mol |
| 3mer binding site 1 | -3.33 kcal/mol |
| 3mer binding site 2 | -3.15 kcal/mol |
| 5mer binding site 1 | -4.8 kcal/mol |
| 5mer binding site 2 | -4.41 kcal/mol |
| 7mer binding site 1 | -4.83 kcal/mol |
| 7mer binding site 2 | -4.63 kcal/mol |

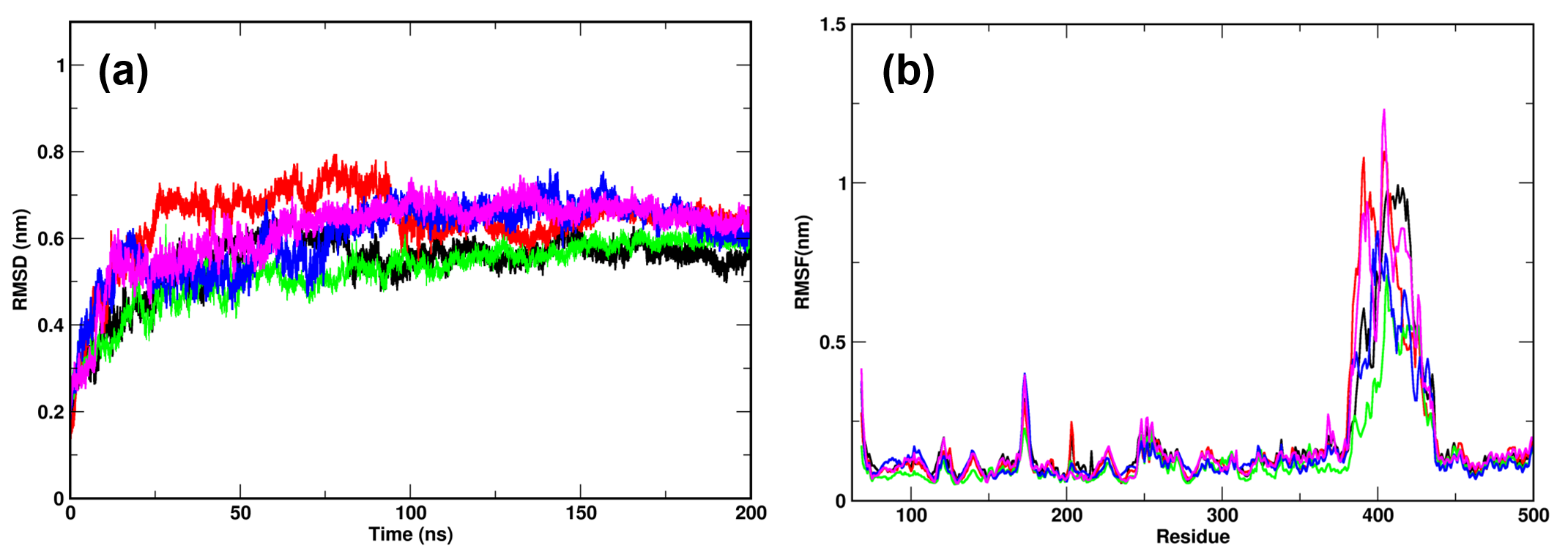

Figure S1. **Structural stability of catalase in single-chain PE-bound complexes.** (a) Backbone Cα RMSD and (b) per-residue RMSF of catalase over 200 ns of MD simulation, shown in triplicate for the 3-, 5-, 7-, and 14-mer single-chain complexes. RMSD traces show rapid initial relaxation followed by equilibration to a stable plateau (~0.5–0.7 nm) after 50–100 ns; RMSF traces identify the wrapping loop (residues 380–430) as the principal region of conformational flexibility across all replicates, irrespective of PE chain length.

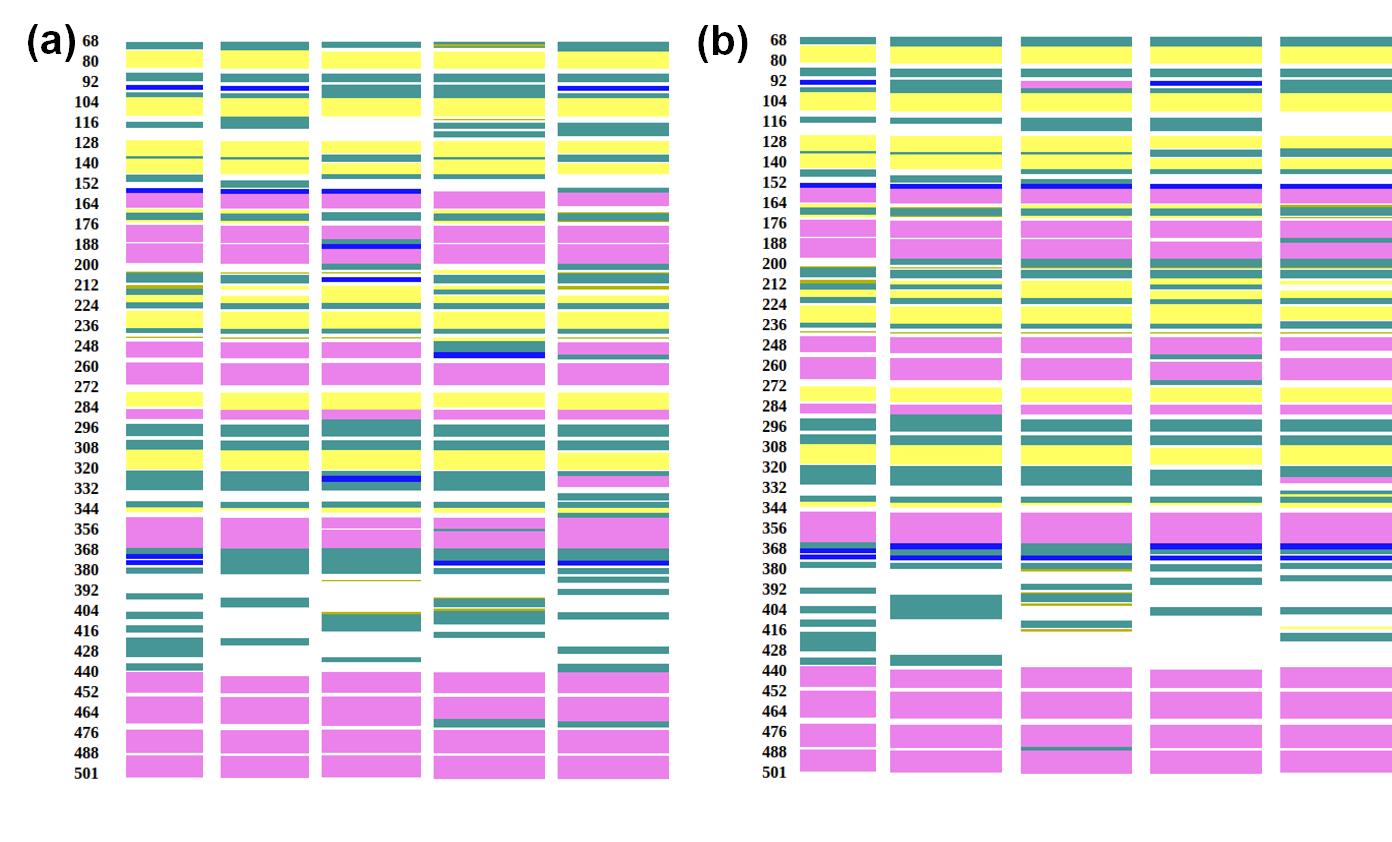

Figure S2. **DSSP-based secondary structure analysis of catalase upon PE binding.** Per-residue secondary structure content (α-helix, β-sheet, turn/coil) of catalase in (a) single-chain PE-bound complexes (1PE3mer, 1PE5mer, 1PE7mer, 1PE14mer) and (b) multichain PE-bound corona complexes, each compared against the unbound wild-type (WT) apo structure over the 200 ns production trajectory.
